# Liver Endothelium Promotes HER3-mediated Cell Survival in *KRAS* Wild-type and Mutant Colorectal Cancer Cells

**DOI:** 10.1101/2021.04.27.441690

**Authors:** Moeez Rathore, Michel’le Wright, Rajat Bhattacharya, Fan Fan, Ali Vaziri-Gohar, Jordan Winter, Zhenghe Wang, Sanford D Markowitz, Joseph Willis, Lee M. Ellis, Rui Wang

## Abstract

We previously showed that liver endothelial cells (ECs) secreted soluble factors in a paracrine fashion and activated human epidermal growth factor receptor 3 (HER3, also known as ERBB3) for promoting colorectal cancer (CRC) growth and chemoresistance. However, RAS proteins play a critical role in receptor tyrosine kinase signaling pathways, and *KRAS* mutations mediate CRC resistance to therapies targeting EGFR, another HER protein. Therefore, the role of *KRAS* mutation status in EC-induced HER3 activation and CRC survival was investigated as it has therapeutic implications. We used CRC cell lines and patient-derived xenografts harboring *KRAS* wild-type or mutant genes and demonstrated that liver EC-secreted factors promoted HER3-mediated CRC cell growth independent of *KRAS* mutation status. Also, blocking HER3 in CRC cells by siRNAs or a HER3 antibody seribantumab blocked EC-induced CRC survival. Our findings highlight the potential of utilizing HER3 targeted therapies for treating patients with mCRC independent of *RAS* mutational status.

## INTRODUCTION

In the United States, colorectal cancer (CRC) is the second-leading cause of cancer-related deaths, with estimated deaths of >50,000 patients per year ^1^. Close to 25% of CRC cases are metastatic (mCRC) at the time of diagnosis and over 20% patients with Stages 1-3 CRC will develop into mCRC. With the exception of immune checkpoint inhibitors for patients with microsatellite instability-high (MSI-H), nearly 50% of patients with mCRC do not respond to systemic therapies and the duration of response is less than one year ^2,3^. As a result, patients with unresectable mCRC have a 5-year survival rate well below 20%. Therefore, a better understanding of the regulations of CRC cell survival is urgently needed for the development of novel therapeutic strategies for patients with mCRC.

As over 80% of patients with mCRC develop metastases in the liver ^4^, our laboratory aims to elucidate potential roles of the liver microenvironment on CRC survival pathways, with a focus on the liver endothelial cells (ECs) as they represent more than 50% of all stromal cells in the liver ^5^. Preclinical studies from other groups in gastric, liver and other types of cancers used established human umbilical vein ECs and showed that ECs secreted soluble factors and activated cancer-promoting signaling pathways (such as AKT, NFκB) in cancer cells ^6-9^, known as angiocrine signaling. In contrast, our laboratory isolated primary ECs from non-neoplastic liver tissues to recapitulate the liver EC microenvironment and demonstrated that liver ECs secreted soluble factors and activated cancer stem cell-associated Notch and Nanog pathways in CRC cells ^10-12^. More recently, we determined that liver EC-secreted factors specifically activated human epidermal growth factor receptor 3 (HER3, also known as ERBB3), a member of the receptor tyrosine kinase (RTK) HER family, and its downstream target AKT, leading to increased cell proliferation and resistance to 5-fluorouracil (5-FU) induced apoptosis in CRC ^13^. In that study, we also determined that EC-secreted factors activated HER3 and AKT independent of EGFR or HER2.

Constitutive activation mutations in *RAS* genes (90% mutations found in *KRAS* ^14^) occur in ∼50% CRC ^15^. Because KRAS is a key mediator of RKT and other pro-survival pathways^16^, mutations in *KRAS* genes lead to high levels of activations in these pathways and patients with mutant *KRAS* mCRC have markedly worse prognosis compared to those with wild-type *KRAS* ^17-19^. As a result, anti-RTK targeted therapies have limited effect in cancer patients with *RAS* mutations ^20^. Specifically, EGFR targeted therapies with antibodies and inhibitor failed to improve the outcomes of patients with *KRAS* mutant mCRC ^21-24^, and *KRAS* mutations are approved by the FDA as resistance markers for CRC response to EGFR targeted therapies. Because both EGFR and HER3 are RTKs in the HER protein family and trigger KRAS activation ^25^, the role of *KRAS* mutations in EC-induced HER3 activation and downstream effects needs to be elucidated. Moreover, the effects of blocking HER3 on CRC cell survival needs to be determined in CRC cells with different *KRAS* mutation status for utilizing HER3-targeted therapies for treating patients with mCRC with different *RAS* mutational status.

In this present study, we first used a patient derived xenograft (PDX) tumor model and determined that conditioned medium (CM) from liver ECs that contains EC-secreted soluble factors increased CRC tumor growth in both *KRAS* wild-type and mutant PDXs. We then used multiple *KRAS* wild-type and mutant CRC cell lines and demonstrated that liver ECs activated the HER3-AKT signaling pathway and increased CRC cells resistant to 5-FU regardless of the *KRAS* mutation status. Moreover, the EC-induced pro-survival effects in CRC cells were completely blocked by HER3 inhibition, either with siRNAs or the HER3 antibody seribantumab. We also used an orthotropic liver injection xenograft tumor model and determined that blocking HER3 with seribantumab decreased CRC tumor growth and sensitized CRC to 5-FU chemotherapy *in vivo*. Our results demonstrated that liver EC-induced HER3 activation and CRC survival were independent of *KRAS* mutation status. As the HER3 antibody seribantumab is already being assessed in a clinical trial for treating cancer patients with NRG-1 gene fusions (NCT04383210), this study highlighted the potential for a rapid translation of treating patients with *KRAS* wild-type and *KRAS* mutant mCRC with the combination of 5-FU based chemotherapies and seribantumab, and potentially other HER3 antibodies.

## RESULTS

### CM from liver ECs promoted growth of CRC PDX tumors with wild-type or mutant *KRAS*

We firstly used a proof-of-principle subcutaneous (subQ) xenograft tumor model to determine the effects of liver ECs on CRC growth. CRC PDX tissues harboring either wild-type (*KRAS* WT) or mutant *KRAS* (*KRAS* mut.) genes were subQ implanted in an inoculation mixture of Matrigel and control CM (CM from HCP-1 CRC cells) or CM from human primary liver ECs (EC-1). To maintain the effects of CM on PDXs during the experiment, CM from HCP-1 cells or EC-1 were subQ injected at the inoculation sites once a week throughout the experiment. Our results showed that, in both *KRAS* WT and *KRAS* mut. PDXs, liver EC CM resulted in over 4-fold increases in tumor growth 2-fold increases in tumor weights compared to CRC CM (Fig. 1).

**Figure 1.**
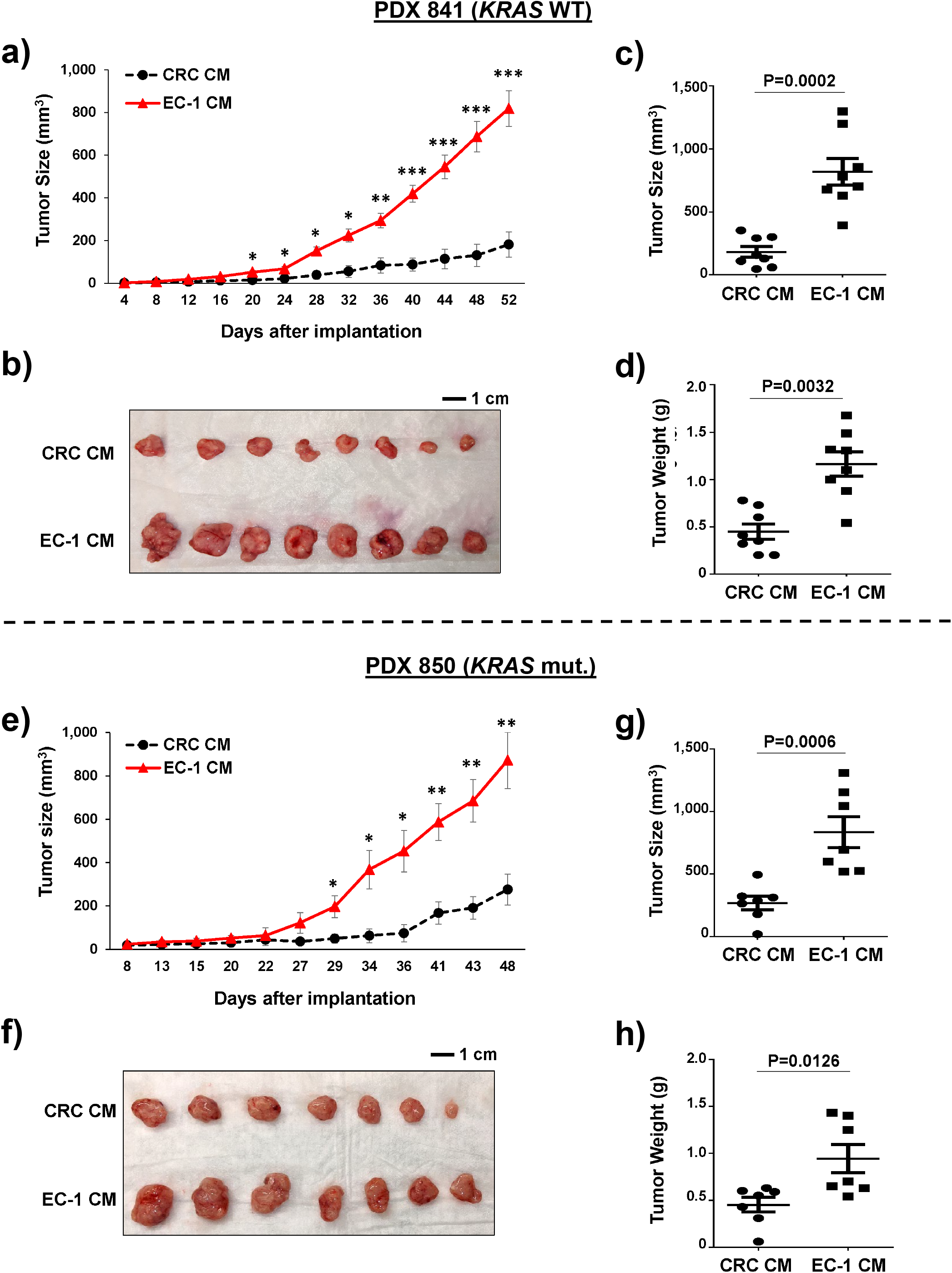
Liver ECs promoted *KRAS* wildtype and mutant CRC PDX tumor growth *in vivo*. CRC PDXs with wild-type (*KRAS* WT) and mutant (*KRAS* mut.) *KRAS* genes were subQ implanted with CM from HCP-1 cells (CRC CM) or primary liver EC (EC-1 CM) (EC-1 CM). **(a, e)** Tumor size measurement over time showed that EC-1 CM promoted PDX tumor growth. Mean -/+ SD, *P<0.01, **P<0.001, ***P<0.0001 Wilcoxon rank-sum test. **(b, f)** Pictures of tumors harvested from each group. Scale bars, 1 cm. **(c, d, g, h)** Scatter plots of tumor sizes and weights after tissue harvest. Mean +/- SD, P value by one-way ANOVA.

### CM from liver ECs activated the HER3-AKT pathway and increased cell survival in CRC cells with different gene mutation profiles

To further elucidate the effects of EC CM on CRC cell growth and involved signaling pathways, we used two primary liver EC lines (EC-1 and EC-6) and multiple CRC cell lines that carried either a wild-type *KRAS* gene (SW48 and Caco2) or mutant *KRAS* (HCP-1). Moreover, we used modified HCT116 and DLD-1 cells that have the *KRAS* mutant alleles being knocked out (*KRAS* mut. KO), therefore only express wild-type *KRAS*, and their paired parental cells with mutant *KRAS* to further determine the specific roles of *KRAS* mutations in liver ECs activating HER3-AKT and promoting cell survival in CRC cells with different *KRAS* mutation status (Supplementary Table 1).

**Table 1.** Gene mutation profiles of CRC cell lines.

| **Cell Line** | ***KRAS*** | ***PIK3CA*** | ***TP53*** | **MSI status** |
| --- | --- | --- | --- | --- |
| **SW48** | **+** | **+** | **+** | Instable |
| **Caco2** | **+** | **+** | E204X | Stable |
| **HCP-1** | G12D/+ | H1047R/+ | NA | Instable |
| **HCT116** | G13D/+ | H1047R/+ | **+** | Instable |
| **HCT116**<br>***KRAS* mut. KO** | **-/+** | H1047R/+ | **+** | Instable |
| **DLD-1** | G13D/+ | E545K/+ | S241F | Instable |
| **DLD-1**<br>***KRAS* mut. KO** | **-/+** | E545K/+ | S241F | Instable |

**+**: wildtype, **-** or **KO**: knockout, **mut.**: mutation, **X**: stop,
**NA**: not available, **MSI**: microsatellite instability

We harvested liver EC CM containing EC-secreted factors and added to CRC cells, with CM from CRC cells themselves as control CM. Compared with control CM, CM from liver ECs dramatically increased the levels of phosphorylation in HER3 and its downstream target AKT, demonstrating activation of the HER3-AKT survival pathway in both *KRAS* wild-type and mutant CRC cells used (Fig. 2). Moreover, we found that the modified HCT116 and DLD-1 cells with wild-type *KRAS* alleles responded to EC CM similar to unmodified HCT116 and DLD-1 parental cells with mutant *KRAS* and also had increased phosphorylation in HER3 and AKT when treated by EC CM (Fig. 2b). These findings suggested that EC CM induced HER3 and AKT activation independent of *KRAS* mutations. We noticed that under the control conditions, HCP-1 cells and modified HCT116 cells with *KRAS* mut. KO had higher basal levels of phosphorylation in HER3 and AKT than other cells. However, other *KRAS* mutant cells (HCT116 and DLD-1) and *KRAS* wild-type cells (SW48, Caco2, and modified DLD-1) had similar non-detectable basal levels of HER3 and AKT phosphorylation. The high basal levels of HER3 and AKT phosphorylation are likely to be cell line-specific and not related to *KRAS* mutation status.

**Figure 2.**
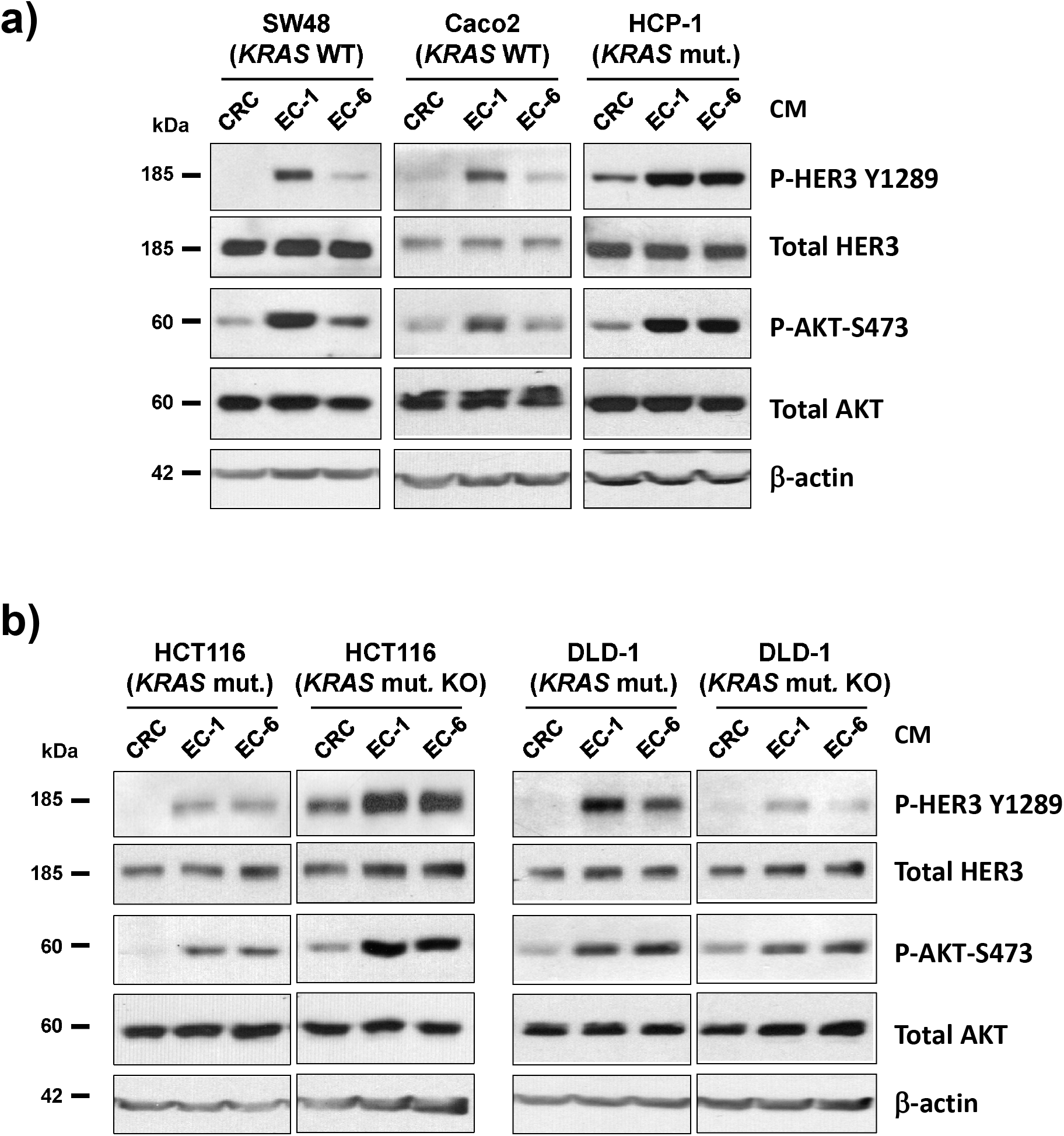
CM from liver ECs activated HER3-AKT in *KRAS* wild-type and mutant CRC cells. CRC cells were incubated with control CRC CM or CM from different primary liver ECs (EC-1 and EC-6) for 30 minutes. Western blotting showed that EC CM increased levels of HER3 and AKT phosphorylation in **(a)** SW48 and Caco2 cells with wild-type *KRAS* (*KRAS* WT) and HCP-1 cells with mutant *KRAS* (*KRAS* mut.), and **(b)** HCT116 and DLD-1 cells with *KRAS* mutant (*KRAS* mut.,) and their sub-clones with the mutant *KRAS* alleles knocked out (*KRAS* mut. KO). Total levels of HER3, AKT, and β-actin were used as loading controls. Data represents results of at least three independent experiments.

We then sought to determine the effects of EC-induced HER3-AKT activation on cancer cell survival, including cell proliferation and response to chemotherapy. CRC cells were incubated in control CM or EC CM and then treated without or with a cytotoxic chemotherapy agent 5-FU. The MTT assay was used to determine cell viability under different conditions and data were presented as percent viability relative to cells in CRC control CM without 5-FU (Fig. 3). We found that in the absence of 5-FU, CM from liver ECs significantly increased cell viability in CRC cells compared to CRC CM. Specifically, CRC cells with *KRAS* mutations had greater levels of increases in cell viability (∼2.7 to 3.5-fold increases in HCP-1, HCT116 and DLD-1 cells) compared to CRC cells with wild-type *KRAS* (<2-fold increases in SW48 and Caco2 cells). Moreover, we compared the EC CM-induced proliferation between HCT116 and DLD-1 parental and modified cells, and found that EC CM caused a 3.5-fold increase in HCT116 parental cells compare to a 2.3-fold increase in HCT116 *KRAS* mut. KO cells, and a 2.8-fold increase in DLD-1parental cells compare to 2.2 fold increase in DLD-1 KRAS mut. KO cells (Fig. 3b, c). These findings confirmed that when stimulated by EC CM, *KRAS* mutant CRC cells had higher cell viability than that of *KRAS* wild-type cells.

**Figure 3.**
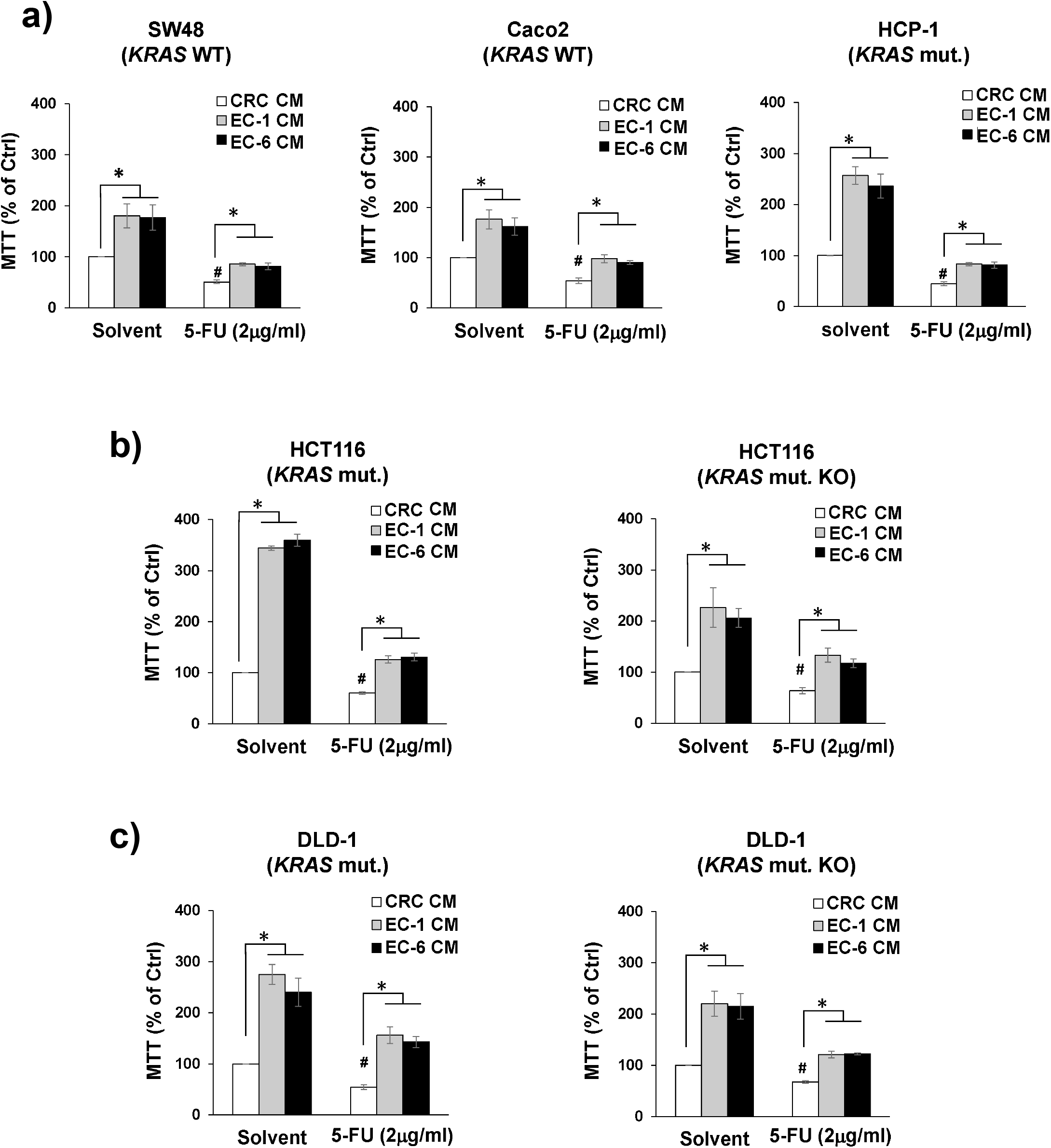
CM from liver ECs increased CRC cell viability and resistance to chemotherapy in *KRAS* wild-type and mutant CRC cells. CRC cells were incubated with control CRC CM or CM from different primary liver ECs (EC-1 and EC-6) and treated without (Solvent) or with 5-FU in CM for 72 hours. The MTT assay showed that CM from liver ECs increased cell viability in **(a)** SW48 and Caco2 cells with wild-type *KRAS* (*KRAS* WT) and HCP-1 cells with mutant *KRAS* (*KRAS* mut.), and **(b, c)** HCT116 and DLD-1 cells with *KRAS* mutant (*KRAS* mut.,) and their sub-clones with the mutant *KRAS* alleles knocked out (*KRAS* mut. KO). Relative cell viability was presented as % of control groups treated with CRC CM and solvent. Mean +/- SEM of at least three experiments, *p<0.01 *t*-test, ^#^p<0.01 *t*-test compared to control groups with CRC CM and solvent.

For cell response to chemotherapy, we treated CRC cells with a clinically relevant dose of 5-FU (2 μg/ml) ^26^, which increased the levels of apoptotic markers such as cleaved PARP and Caspase 3 and led to sufficient apoptosis in CRC cells in our previous studies ^10,11,13^. We confirmed that when treated with CRC CM, 5-FU was effective and decreased CRC cell viability to ∼50% relative to control groups without 5-FU. In contrast, 5-FU had limited effects on CRC cells in the presence of EC CM, leading to 100%-150% relative cell viability in both *KRAS* wild-type and mutant cells. These findings suggested that EC CM blocked the cytotoxic effects of 5-FU in CRC cells regardless of the *KRAS* mutation status and CRC cells became more resistant to 5-FU when incubated in EC CM.

### HER3 mediated EC CM-induced AKT activation and cell survival in CRC cells with different gene mutation profiles

To determine the role of HER3 in mediating EC CM-induced AKT activation and CRC cell survival, we firstly used siRNAs to knock down HER3 expressions in CRC cells. We found that EC CM could no longer induce AKT phosphorylation in CRC cells without HER3 expression in both *KRAS* wild-type and mutant CRC cells (Fig. 4). This observation was further validated by blocking HER3 with a humanized HER3-specific antibody seribantumab, which demonstrated significant HER3 inhibition in previous studies ^27,28^. We showed that seribantumab completely blocked EC-induced HER3 and AKT phosphorylation in all cell lines we used (Fig. 5). Taken together, these findings demonstrated that EC-induced AKT activation was mediated by HER3 and HER3 inhibition completely blocked EC CM-induced AKT activation in CRC cells independent of the *KRAS* mutation status. As we previously demonstrated that EC activated HER3 and AKT independent of EGFR and HER2 in CRC cells ^13^, we confirmed that EC CM did not activate EGFR and HER2, and blocking EGFR or HER2 did not affected EC-induced HER3 and AKT activation in all CRC cell lines used in these studies (data not shown).

**Figure 4.**
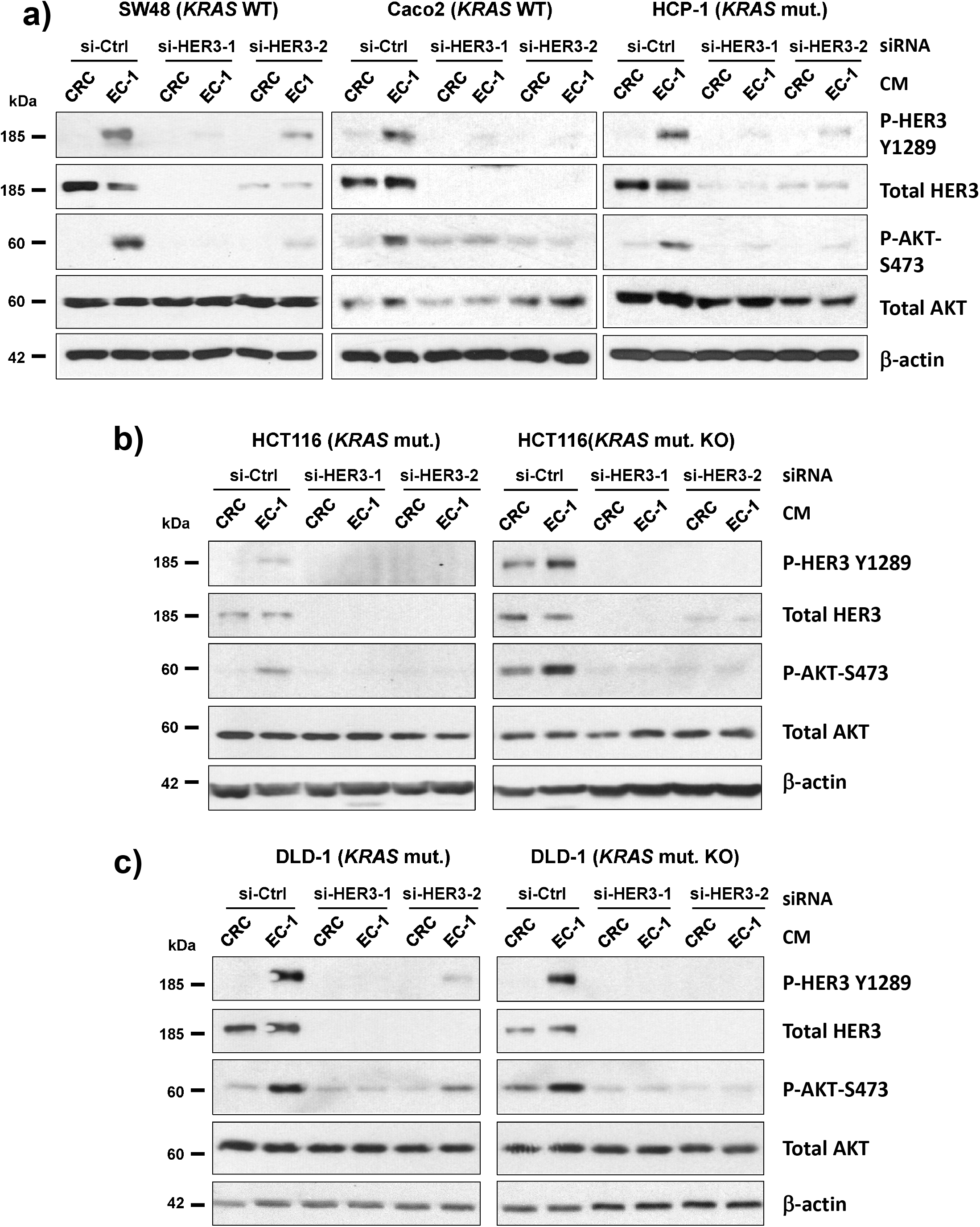
HER3 knockdown by siRNAs inhibited liver EC-induced HER3-AKT activation in *KRAS* wild-type and mutant CRC cells. CRC cells were transfected with control (si-Ctrl) or HER3-specific siRNAs (si-HER3-1 and si-HER3-2) and then incubated with control CRC CM or CM from different primary liver ECs (EC-1 and EC-6). The Western blotting showed that HER3 siRNAs decreased HER3 protein levels, and blocked liver EC CM-induced AKT phosphorylation in **(a)** SW48 and Caco2 cells with wild-type *KRAS* (*KRAS* WT) and HCP-1 cells with mutant *KRAS* (*KRAS* mut.), and **(b, c)** HCT116 and DLD-1 cells with *KRAS* mutant (*KRAS* mut.,) and their sub-clones with the mutant *KRAS* alleles knocked out (*KRAS* mut. KO). Total levels of HER3, ATK and β-actin were used as loading controls. Data represents results of at least 3 independent experiments.

**Figure 5.**
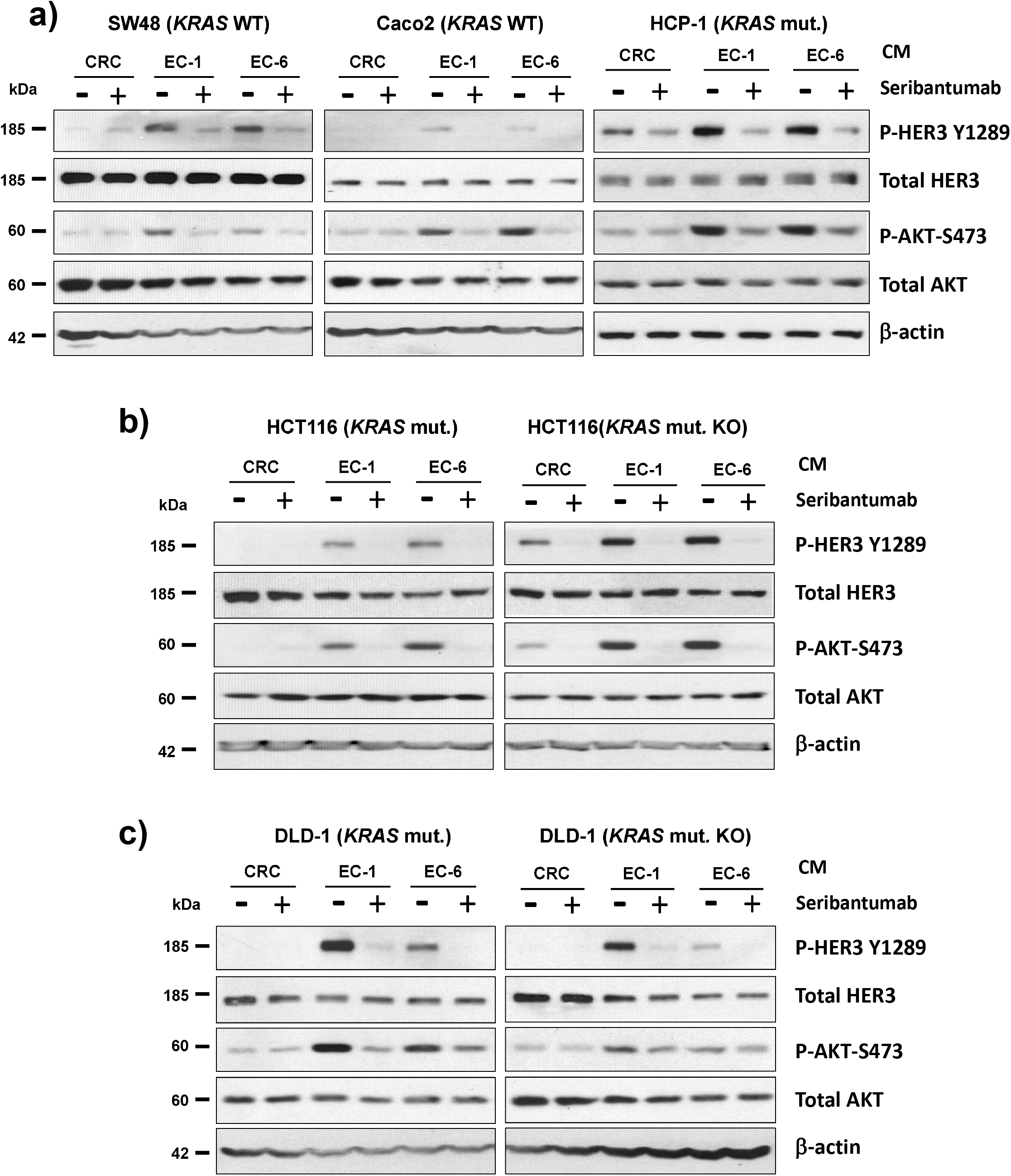
HER3 antibody seribantumab blocked liver EC-induced HER3-AKT activation in *KRAS* wild-type and mutant CRC cells. CRC cells were incubated in control CRC CM or CM from different primary liver ECs (EC-1 and EC-6) either in presence or absence of the HER3 antibody seribantumab (200 μg/ml) for 30 minutes. The Western blotting showed that seribantumab blocked liver EC CM-induced HER3 and AKT phosphorylation in **(a)** SW48 and Caco2 cells with wild-type *KRAS* (*KRAS* WT) and HCP-1 cells with mutant *KRAS* (*KRAS* mut.), and **(b, c)** HCT116 and DLD-1 cells with *KRAS* mutant (*KRAS* mut.,) and their sub-clones with the mutant *KRAS* alleles knocked out (*KRAS* mut. KO). Total levels of HER3, ATK and β-actin were used as loading controls. Data represents results of at least 3 independent experiments.

We then used the MTT assay to determine the effects of blocking HER3 on EC-induced CRC cell proliferation and resistance to 5-FU induced apoptosis. First, we confirmed that seribantumab significantly blocked EC-induced proliferation in CRC cells with different mutation profiles (Suppl. Fig. 1). Subsequently, we incubated CRC cells in CM with 5-FU either alone or in combination with seribantumab (Fig. 6). EC CM significantly increased cell viability and seribantumab blocked EC-induced proliferation, as expected, in all CRC cells used in these studies. Also, we confirmed that 5-FU decreased cell viability in all conditions but was less effective when treated in the presence of EC CM as a result of EC-induced chemoresistance. In contrast, when we treated CRC cells with both seribantumab and 5-FU, EC-induced resistance to 5-FU was completed blocked by seribantumab and CRC cells had significantly lower levels of cell viabilities compared to those from the groups treated by either 5-FU or seribantumab alone. Similar results were found in HCT116 and DLD-1 CRC cells with or without *KRAS* mutant alleles (Fig. 6b, c) and confirmed that the effects of HER3 inhibition on EC-induced cell proliferation and resistance to chemotherapy were independent of *KRAS* mutations. Taken together, these findings demonstrated that HER3 inhibition blocked EC-induced CRC cell proliferation and chemoresistance, resulting sensitizing CRC cells to 5-FU treatment in both *KRAS* wild-type and mutant CRC cells.

**Figure 6.**
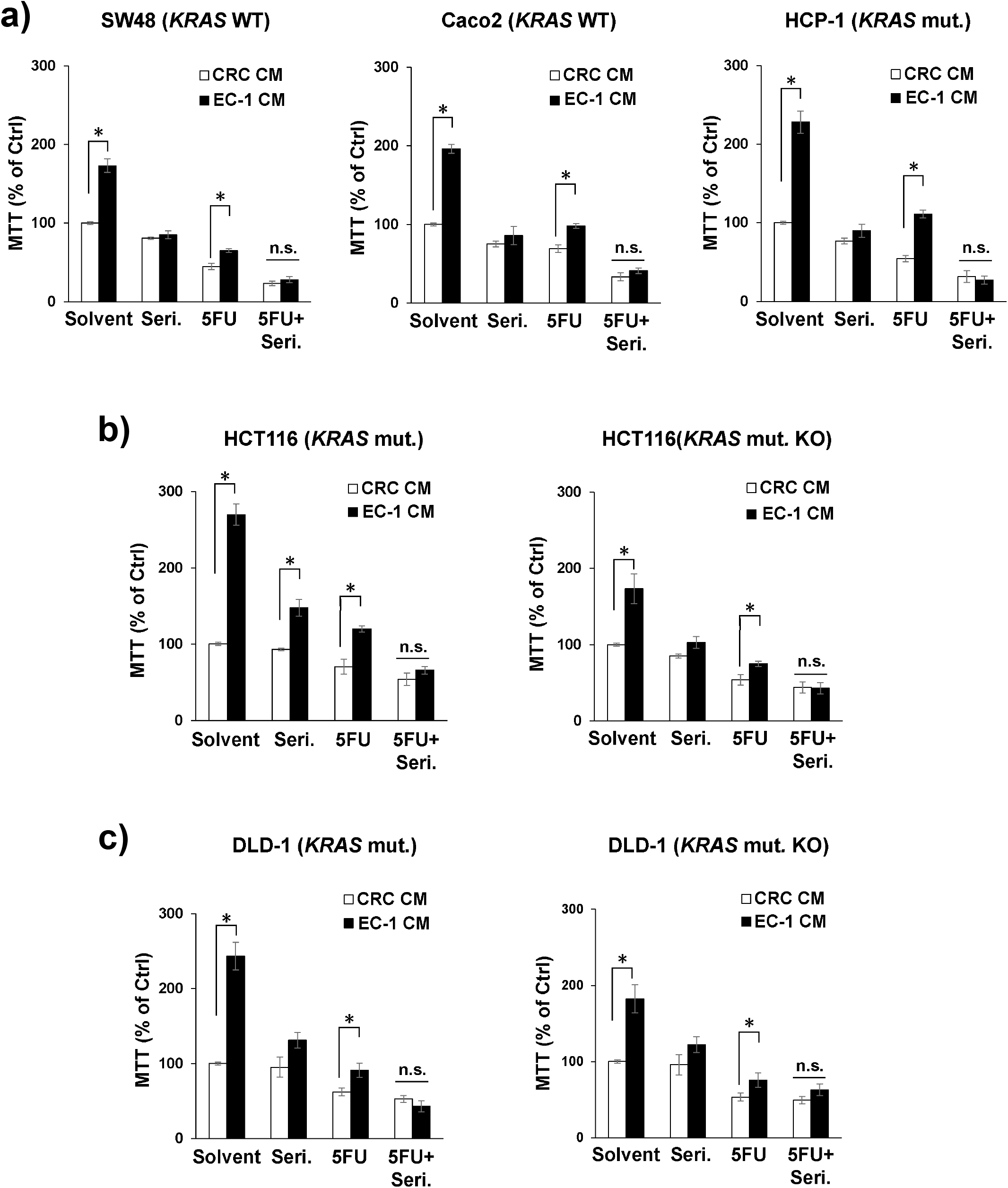
HER3 antibody seribantumab blocked liver EC-induced CRC cell viability and chemoresistance in *KRAS* wild-type and mutant CRC cells. CRC cells were treated with control CM (CRC) or CM from primary liver ECs (EC-1) and with the HER3 antibody seribantumab (Seri., 200 μg/ml) or 5-FU for 72 hours. The MTT assay showed that seribantumab or 5-FU decreased EC CM-induced CRC cell viability and the combination of seribantumab and 5-FU caused the lowest cell viability in **(a)** SW48 and Caco2 cells with wild-type *KRAS* (*KRAS* WT) and HCP-1 cells with mutant *KRAS* (*KRAS* mut.), and **(b, c)** HCT116 and DLD-1 cells with *KRAS* mutant (*KRAS* mut.,) and their sub-clones with the mutant *KRAS* alleles knocked out (*KRAS* mut. KO). Relative cell viability was presented as % of control groups treated with CRC CM and solvent. Mean +/- SEM. *p<0.01 *t*-test. n.s., not significantly different (p>0.05) by *t*-test.

### HER3 inhibition sensitized CRC tumors in the liver to 5-FU *in vivo*

As anti-EGFR targeted therapies demonstrated significant anti-cancer effects only in patients with *KRAS* wild-type mCRC, it is necessary to develop targeted therapies that are also effective in patient with *KRAS* mutant mCRC. Therefore, we focused on assessing the possibility of treating patients with *KRAS* mutant mCRC with HER3 targeted therapies. To do so, we used a liver injection orthotopic xenograft model to determine the effects of blocking HER3 with the HER3 antibody seribantumab on CRC tumors in the liver. We firstly isolated murine primary liver ECs from athymic nude mice and generated murine EC CM to confirm that secreted factors from murine liver ECs activated HER3 and AKT in human CRC and promoted cancer cell growth (Suppl. Fig. 2). We then injected luciferase-labeled human CRC cells (HCP-1, *KRAS* mutant) into the livers of athymic nude mice to recapitulate CRC liver metastases developed in patients with mCRC. After confirming tumor burden, mice were randomized and then treated with seribantumab (20 mg/kg) either alone or in combination with 5-FU (20 mg/kg) (Fig. 7). Compared to the control group, monotherapy treatments with either seribantumab or 5-FU caused significant but modest decreases in tumor growth as determined by tumor burden over time. In contrast, combination of seribantumab and 5-FU dramatically decreased tumor growth, leading to over 70% decrease in tumor burden (Fig. 7a) and ∼50% decrease in weights of tumor-bearing livers (Fig. 7b) compared to those of the control group. However, we noticed that seribantumab or 5-FU monotherapy treatments did not cause significantly changes in weights of tumor-bearing livers. This can be partially explained as the experiment ended 21 days after implanting CRC cells due to fast tumor growth. As a result, tumor-bearing mice only received four drug administrations, which could significantly affect the effects of seribantumab and 5-FU on CRC tumors. Also, the tumor-bearing livers were measured with residual non-neoplastic normal liver tissues. As shown in Fig. 7c with the livers harvested from all four groups and an enlarged image of two livers from the control group as an example, significant portion of the liver was occupied by tumors derived from implanted CRC cells (T, pale/light red tissues) left with a small fraction of non-neoplastic normal liver (N, dark red tissues). Compared to the control group, nearly half of the animals from 5-FU or seribantumab treatment groups had noticeable portions of normal liver tissues remained. In addition, bioluminescence measurements on Day 21 immediate prior tissue harvesting showed that tumor burdens from monotherapy treatment groups were significantly lower than those from the control group (Fig. 7a). Taken together, our results showed that blocking HER3 with seribantumab was able to decrease CRC tumor growth in the liver, and more importantly, the combination of seribantumab and 5-FU decreased tumor growth even further compared to monotherapies. These findings suggested that blocking HER3 sensitized CRC tumors to 5-FU and increased the anti-tumor effects of a relatively low dose of 5-FU on CRC tumors established in the liver.

**Figure 7.**
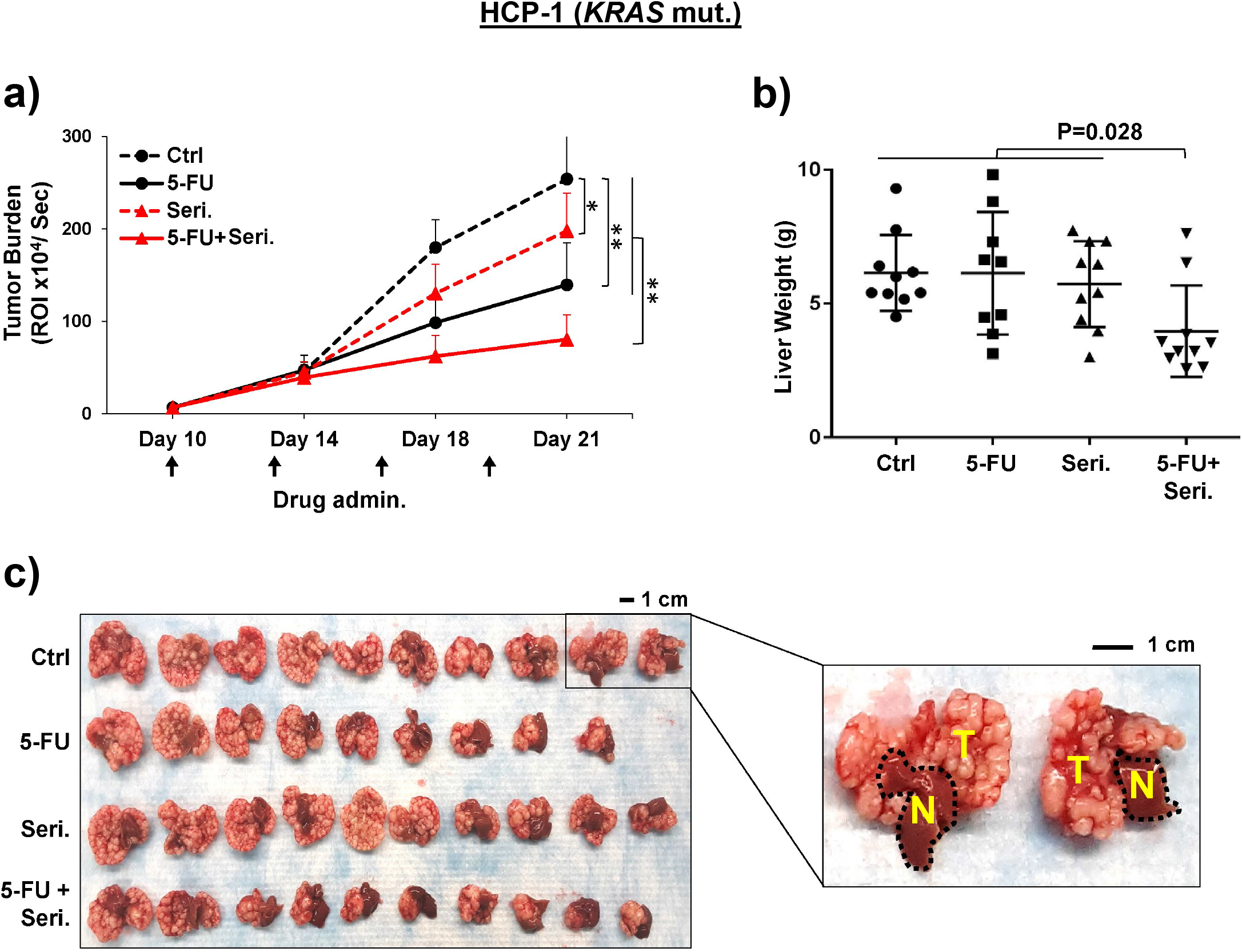
HER3 antibody seribantumab sensitized *KRAS* mutant CRC tumors to chemotherapy *in vivo*. Luciferase reporter-labeled HCP-1 CRC cells (*KRAS* mut.) were injected into the livers of athymic nude mice. Once tumor burden confirmed by bioluminescence 10 days after injection (Day 10), mice were treated with control vehicle (Ctrl), 5-FU alone, seribantumab alone (Seri.), or the combination of 5FU and seribantumab (5FU+Seri.) in every three days (black arrow). **(a)** Tumor burdens measured by the luminescence IVIS system over time. Mean -/+ SD, *P<0.02, **P<0.001 Wilcoxon rank-sum test between groups on Days 18 and 21. **(b)** Scatter plots of weighs of the tumor-bearing livers. Mean +/- SD, P value by one-way ANOVA. **(c)** A picture of the tumor-bearing livers harvested from each group (left), and an enlarged image of two samples with tumor tissues (T, pale light red regions) and non-neoplastic normal liver tissues (N, dark red regions). Scale bars, 1 cm.

## DISCUSSION

Previous studies have demonstrated critical roles of the stromal microenvironment in cancer cells in different types of cancer ^29-32^. Studies from our group and others suggested that ECs, an understudied component of the microenvironment, also have pro-survival roles in CRC and other types of cancer ^10,12,13,33^. As Liver is the most common site of CRC distant metastases and it has a unique EC-rich microenvironment, the present study sought to elucidate the role of liver ECs in mediating CRC cell survival. Moreover, as existing targeted therapies have little effect on patients with *KRAS* mutant mCRC ^34,35^, we directly compared the effects of liver ECs among CRC cells with different *KRAS* mutation statuses. We determined that liver ECs promoted CRC cell proliferation and chemoresistance by activating cancer cell-associated HER3-AKT in both *KRAS* wild-type and mutant CRC cells. Moreover, we showed that inhibiting HER3 with a HER3 antibody blocked liver EC-induced CRC cell proliferation and chemoresistance in *in vitro* and *in vivo* preclinical models.

To date, neuregulin family proteins have been the only identified ligands for HER3. Because HER3 has an intracellular domain with weak kinase activity and is considered as a “kinase-dead” receptor ^36^, neuregulin-binding leads to HER3 dimerization with HER2 and, to a lesser extent, other HER family receptors to activate downstream pathways including AKT ^25,37^. The canonical activation of HER3-AKT in cancer cells involves PI3K and/or MAPK pathways ^38,39^. When gain-of-function mutations occur in *KRAS* and *PIK3CA* genes, PI3K and MAPK pathways are highly active and are not expected to be further activated by additional stimulations. To our surprise, liver EC-secreted factors activated HER3-AKT in CRC cells independent of mutations in *KRAS* and *PIK3CA* genes. It is possible that the HER3-AKT activation we observed was mediated by other signaling pathways. Indeed, our previous studies demonstrated that EC-induced HER3-AKT activation in CRC cells was independent of HER3 dimerization with HER2 or EGFR and was independent of the HER3 ligand neuregulins ^13^. These findings strongly suggest that EC-secreted factors activate HER3 via a mechanism that has not reported before. Our laboratory is currently identifying the key EC-secreted factor for binding and activating HER3 and elucidating the mechanism of liver EC-induced HER3 activation, which are beyond the scope of this study.

In addition to *KRAS* and *PIK3CA* mutations, CRC cells used in this study harbored mutations in *TP53*. Preclinical studies suggested that there are correlations between resistant to therapies and *TP53* alterations in mCRC ^40-42^. Our results showed that liver ECs induced HER3-AKT activation and cell survival in all cell lines we used, which harbor either wild-type or mutant *TP53*. It suggests that the pro-survival effects of liver ECs are also independent of *TP53* mutations and, more importantly, that HER3-targeted therapy can potentially be used for treating patients with mCRC with different *TP53* alterations.

Previous studies have characterized the pro-survival role of HER3 primarily in breast, ovarian and other cancer types ^43^. In CRC, ∼75% of primary and metastatic tumors express HER3 ^44,45^. cBioPortal analysis of TCGA and multiple other databases showed that majority of the tumors (>94%) do not carry mutation, duplication, or other alterations in HER3. Meanwhile, HER3 overexpression occurs frequently and is associated with poor prognosis in patients with CRC ^46,47^. As a result, HER3 inhibition has been proposed as a promising targeted therapy strategy in different types of cancers.

However, previous clinical studies showed that blocking HER3 had limited anti-tumor effect in several types of cancer by using either HER3 antibodies/inhibitors as monotherapies or in combination with EGFR- or HER2-targeted therapies such as cexucimab and trastuzumab ^48-50^. The HER3 antibody used in this study, seribantumab, has been used in clinical studies for treating breast, lung, and ovarian cancer either alone or in combination with paclitaxel or EGFR/HER2 targeted therapies ^51-53^. These studies showed that seribantumab was only effective in patients with NRG-1 gene fusion, which led to the ongoing CRESTONE trial (NCT04383210) for treating cancer patients with NRG-1 fusions that represent less than 0.2% of all cancer cases. On the other hand, the effects of HER3 antibodies on CRC remain unknown, and there is little work has been done in combining HER3 antibodies with cytotoxic chemotherapy agents that are commonly used in CRC, such as 5-FU. A phase I study with 20 *KRAS* wild-type CRC patients used seribantumab and irinotecan but the combination did not improve patient outcomes (NCT01451632) ^54^. Considering the limited number of patients in this study and irinotecan is rarely being used as a single agent chemotherapy in CRC, the effects of combining seribantumab and cytotoxic chemotherapy on CRC needs to be further determined. Moreover, the effects of HER3 antibodies on *KRAS* mutant CRC remains unknown. Unexpectedly, most clinical trials for HER3 targeted therapies did not assess the levels of expression or phosphorylation of HER3 in the patients, which can significantly affect patient response to HER3 targeted therapies. In July 2020, Daiichi Sankyo, Inc. initiated a clinical trial to use an antibody-drug conjugate agent with a HER3 antibody and an irinotecan derivative compound for treating mCRC patients with HER3 expression (NCT04479436). Although there is no data available from this study yet, it highlighted the needs of developing additional HER3 targeted therapies and a potential of combining HER3 antibodies with cytotoxic chemotherapies for treating patients with HER3 positive mCRC.

In summary, our findings determined a role of liver EC microenvironment in activating HER3 and promoting CRC cell survival independent of *KRAS* mutations and other key oncogenic mutations. With the HER3 antibody seribantumab, we showed that blocking HER3 significantly decreased EC-induced CRC proliferation and sensitized CRC to 5-FU chemotherapy. This work demonstrated a potential therapeutic strategy of using HER3 antibodies/inhibitors in combination with established chemotherapy for treating patients with either wild-type or mutant *KRAS* mCRC.

## METHODS

### Cell lines and tissue culturing

The established CRC cell lines SW48 and Caco2 were purchased from ATCC (Manassas, VA, USA). Parental control and *KRAS* mutant allele-deleted HCT116 and DLD-1 cells were described previously ^55^.The human CRC primary cell line (HCP-1) and human liver parenchymal primary ECs (EC-1 and EC-6, also known as LPEC-1 and -6 in previous studies) lines were isolated and established in our laboratory using MACS microbead-conjugated antibodies and separation columns (Miltenyi Biotec, Bergisch Gladbach, Germany) ^10-13^. Anti-EpCAM was used for CRC cells and ant-CD31 was used for ECs. All CRC cells were cultured in MEM (Sigma-aldrich, St. Louis, MO, USA) supplemented with 5% FBS (Atlanta Biologicals, Atlanta, GA), vitamins (1x), nonessential amino acids (1x), penicillin-streptomycin antibiotics (1 x), sodium pyruvate (1x), and L-glutamine (1x), all from Thermo Fisher Scientific/Gibco (Grand Island, NY, USA). Human liver primary ECs were cultured in Endothelial Cell Growth Medium MV2 (PromoCell, Heidelberg, Germany) supplemented with 10% human serum (Atlanta Biologicals) and antibiotics-antimycotic (1x, Thermo Fisher Scientific/Gibco). Cell line mutation status were determined in previous studies^12,56-59^, and from available databases including Cancer Cell Line Encyclopedia (CCLE) and Catalogue of Somatic Mutations in Cancer (COSMIC).

Murine liver primary ECs were isolated from athymic nude mice also using MACS microbead-conjugated antibodies for murine CD31 (Miltenyi Biotec) and cultured in MV2 EC culture medium as described above. HCP-1 cells, human and murine liver ECs were used within 10 passages, with approximately 1 week per passage. Authentication for all cell lines were done in every 6 months by short tandem repeat (STR) tests. For primary cell lines (HCP-1 and ECs) established in our laboratory, genomic DNA from the original tissues was used for authentication. For cell lines from the ATCC, STR profiles of cell lines cultured in our laboratory were compared with available STR profile database for established cell lines. All cell lines were tested for mycoplasma contamination for every 6 months.

### Reagents

The fully humanized IgG2 anti-HER3 antibody seribantumab (previously known as MM-121) was provided by Merrimack Pharmaceuticals (Cambridge, MA, USA), and is now owned by Elevation Oncology Inc. (New York, NY, USA). The control human IgG antibody for *in vitro* and *in vivo* studies was from Invitrogen (Carlsbad, CA, USA).

Pharmaceutical grade 5-fluorouracil (5-FU) was obtained from the pharmacy at The University of Texas MD Anderson Cancer Center. For all *in vitro* studies, 200 μg/ml seribantumab and 2 μg/ml 5-FU were used. Human *ERBB3* (HER3) specific siRNAs (si-HER3-1: 5’-GCUGAGAACCAAUACCAGA, si-HER3-2: 5’-CCAAGGGCCCAAUCUACAA) and a validated control siRNA were obtained from Sigma-Aldrich.

### Conditioned medium (CM)

0.3×10^6^ of CRC cells or ECs were seeded in T25 culture flasks overnight. The next day, cells were washed two times with 1X PBS and then cultured in 3ml growth medium with 1% FBS (0.1×10^6^ cells/ml) for 48 hours. CM was harvested and centrifuged at 4,000 *g* for 10 minutes to remove cell debris. CM from each CRC cell line was used as controls.

### Western blotting

Cell lysates were processed and run by SDS-PAGE gel electrophoresis as described previously ^11,60^. A HRP conjugated β-actin antibody was obtained from Santa Cruz Biotechnology (Santa Cruz, CA, USA). All other antibodies were from Cell Signaling Technology (Beverly, MA, USA). For each experiment, protein lysates were loaded into two gels and processed at the same time for separate probing for antibodies specific to phosphorylated proteins and total proteins. All membranes were probed with β-actin as a loading control and a representative image was shown for each experiment. Each Western blotting figure shows representative results of at least three independent experiments.

### siRNA transfection

For each transfection, 1 × 10^6^ CRC cells were transiently transfected with 400 pmol siRNAs via electroporation using the Neon Transfection System (Invitrogen, Carlsbad, CA, USA) with 3 pulses of 10 msec at 1,600 V according to the manufacturer’s instructions. Cells were recovered in 5% FBS for 24-48 hours, cultured in 1% FBS overnight, and then incubated in CM for 30 minutes for Western blotting, or up to 72 hours for the MTT assay.

### MTT assay

CRC cells were seeded at 3,000 cells/well in 96-well plates, cultured in 1% FBS overnight and then incubated in CM for 72 hours. When seribantumab (200 μg/ml) or 5-FU (2 μg/ml) was used, cells were pretreated with seribantumab in 1% FBS medium for 6 hours, and then cultured with or without 5-FU and seribantumab in CM for 72 hours. Cell viability was assessed by adding MTT substrate (0.25% in PBS, Sigma-Aldrich) in growth medium (1:5 dilution) for 1 hour at 37 °C. Cells were washed with 1x PBS and then added with 50 μl DMSO. The optical density of converted substrate was measured at 570 nm, and relative MTT was presented as percent of control groups with cells treated with CRC CM only.

### Xenograft tumor models

CRC PDXs harboring wild-type or *KRAS* mutant genes were established previously ^61^. Frozen PDX tumors were expanded in athymic nude mice, sliced into ∼5mm^3^ pieces, and then implanted subcutaneously (subQ) into the right flanks of athymic nude mice in an inoculation matrix (100 μl of 1:1 mix of growth-factor-reduced Matrigel and concentrated HCP-1 or EC-1 CM). After implantation, mice were treated with concentrated CM by subQ injection adjacent to implanted tumors once a week. Tumor volumes were measured with a caliper.

For liver injection orthotropic xenografts, CMV-driven luciferase reporter-labeled HCP-1 CRC cells were suspended in an inoculation matrix (1:1 mix of growth factor-reduced Matrigel and serum-free MEM medium) and injected into the left lobe of the livers in athymic nude mice (1×10^6^ cells in 50 μl/injection). After injection, tumor burden was assessed by bioluminescence with the *In Vivo* Imaging System (IVIS) and D-Luciferin substrate (Xenogen, Alameda, CA, USA) according to the manufacturer’s instructions. When tumor burden was confirmed (Day 10), mice were randomized into four groups with equal tumor burden (n=10/group) and were treated with control IgG (20 mg/kg), 5-FU (20 mg/kg), or seribantumab (20 mg/kg) in 100 μl saline by intraperitoneal (I.P.) injection in every three days on Days 11, 14, 17, and 20. All mice were euthanized when three mice in any group became moribund or their tumor sizes reached 1,000mm^3^. The tumor-bearing livers were harvested for imaging and weighing.

### Statistical analysis

For *in vitro* assays, all quantitative data were reproduced in at least three independent experiments, with multiple measures in each replicate. Groups were compared by two-tailed Student’s *t*-test and data was expressed as means -/+ standard error of the mean (SEM) with significance of P<0.05. For *in vivo* assays, Wilcoxon rank-sum test was used for tumor volume and burden change over time, and one-way ANOVA was used for comparing liver weights between groups. Data was expressed as means -/+ standard deviation (SD) with significance of P<0.05.

## ACKNOWLEDGMENTS

This project was supported by the DOD grant CA171038 (L.M.E.), NIH grants CA225756 (R.W.), P30CA016672 (MDACC CCSG), and P30CA043703 (CWRU CCSG)

## AUTHOR CONTRIBUTIONS

Moeez Rathore: collection and assembly of data, data analysis and interpretation

Rajat Bhattacharya: Data analysis and interpretation

Fan Fan: Collection and assembly of data

Michel’le: Wright collection and assembly of data

Ali Vaziri-Gohar: Data interpretation

Jordan Winter: Data interpretation, manuscript editing

Zhenghe Wang: Resource sharing, data interpretation

Sanford D.: Markowitz Resource sharing, data interpretation

Joseph Willis: Resource sharing

Lee M. Ellis: Conception and design, data analysis and interpretation, financial support,

Rui Wang: Conception and design, collection and assembly of data, data analysis and interpretation, manuscript writing, financial support, final approval of manuscript

## Supplementary Information

**Supplementary Figure 1.**
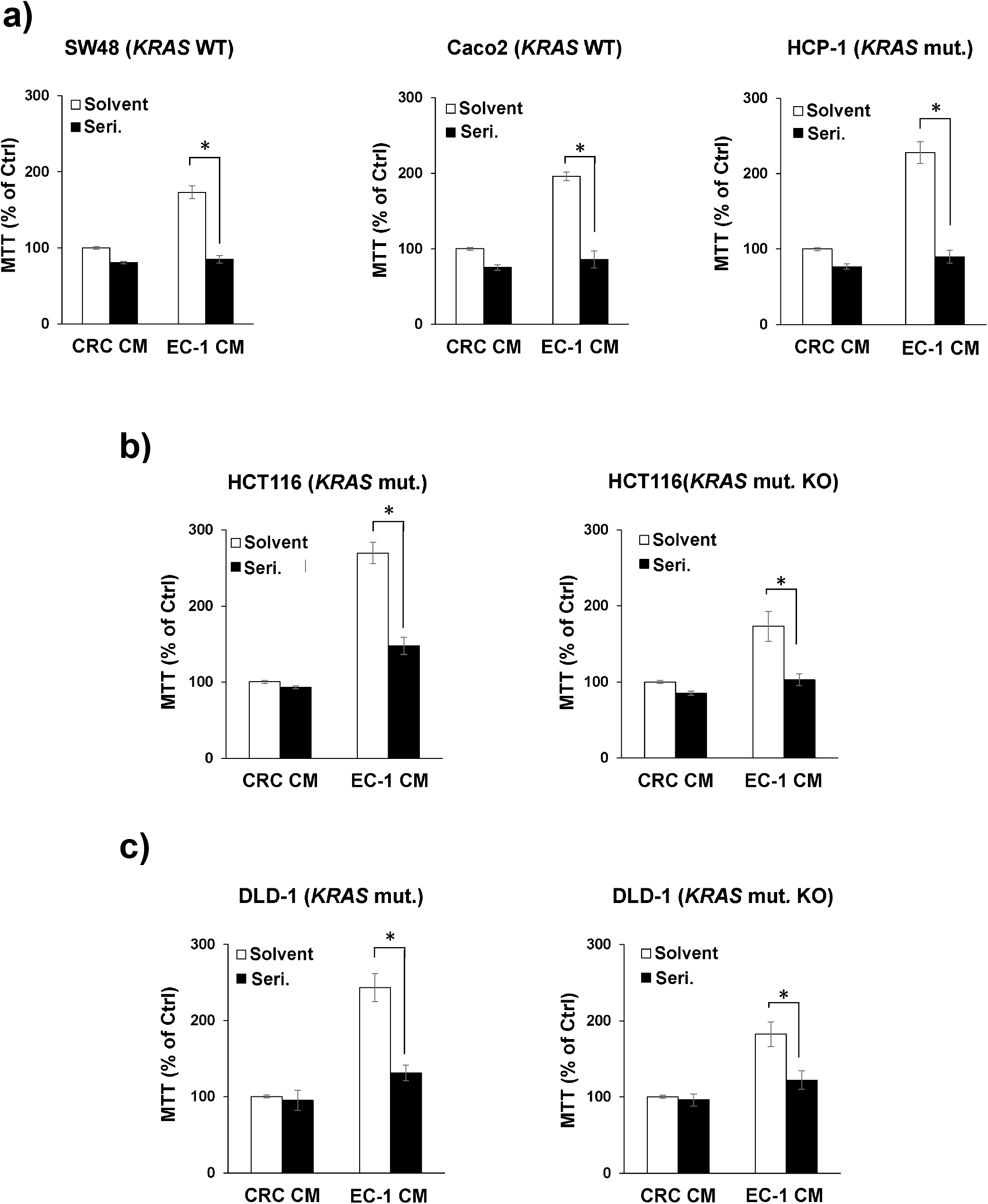
HER3 antibody seribantumab blocked liver EC-induced viability of CRC cells with different mutation profiles. CRC cells with wild-type *KRAS* (SW48 and Caco2), mutant *KRAS* (HCP-1, HCT116, and DLD-1), and sub-clones with the mutant *KRAS* allele knocked out (*KRAS* mut. KO) where treated without or with the HER3 antibody seribantumab (Seri.) in control CM (CRC CM) or primary liver ECs CM (EC-1 CM) for 72 hours. The MTT assay showed that seribantumab significantly blocked EC CM-induced cell viability in CRC cells. Meanb +/- SEM of at least three experiments, relative cell viability was presented as % of control group with CRC CM and solvent. *p<0.01 *t*-test.

**Supplementary Figure 2.**
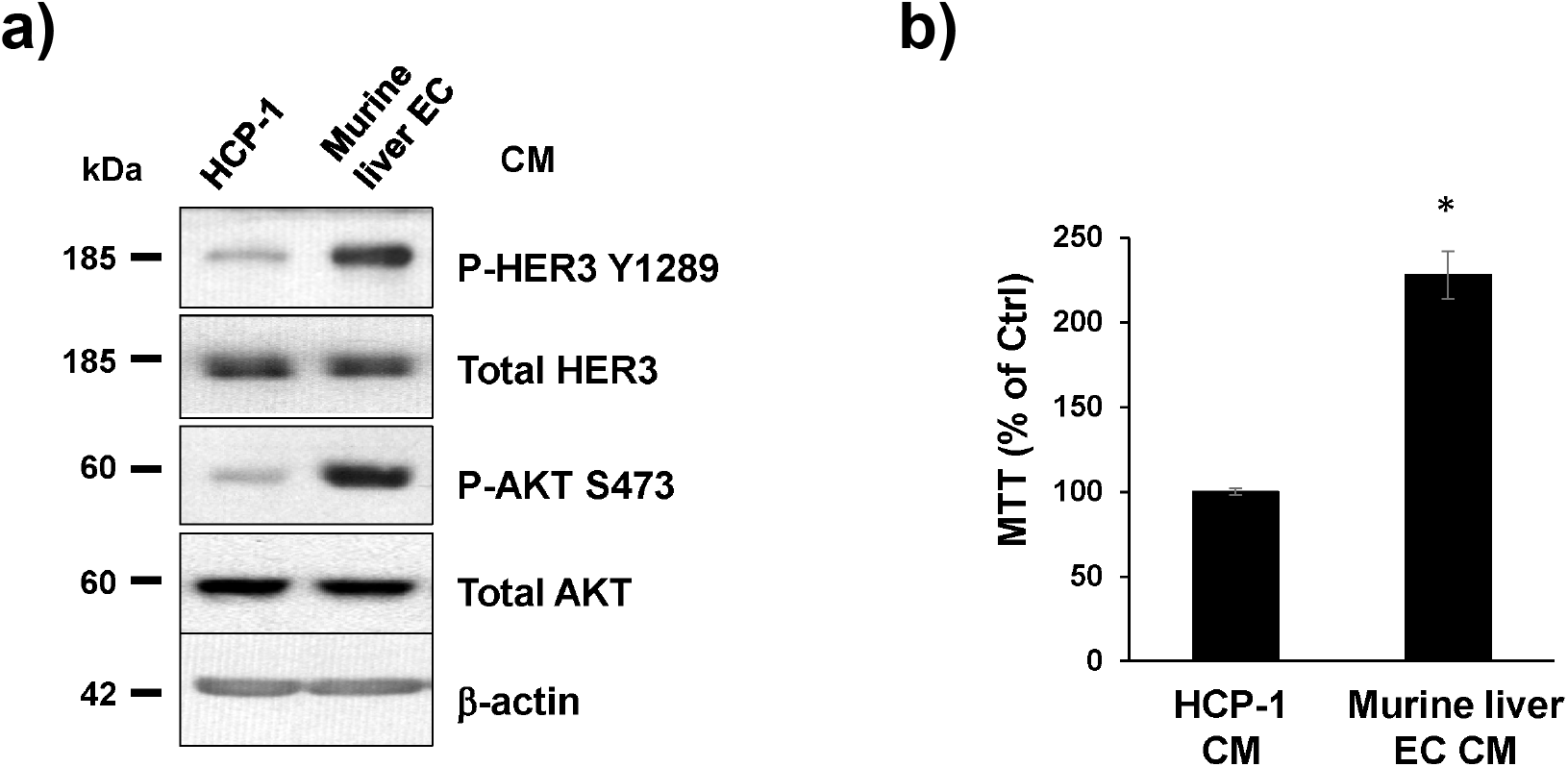
CM from murine primary liver ECs activated HER3-AKT and induced cell growth in human CRC cells. Murine primary liver ECs were isolated from athymic nude mice and cultured for making CM. HCP-1 human CRC cells were incubated in its own CM or in murine liver EC CM. **(a)** Western blotting showed HER3 and AKT phosphorylation in human CRC cells were induced by murine EC CM after 30-minute treatment. The total levels of HER3, ATK and β-actin were used as loading controls. Data represents results of at least three independent experiments. **(b)** The MTT assay showed that CM from murine liver ECs increased human CRC cell proliferation after 72-hour treatment. *p<0.05 *t*-test.

